# Myosin II activity is not required for *Drosophila* tracheal branching morphogenesis

**DOI:** 10.1101/098814

**Authors:** Amanda Ochoa-Espinosa, Stefan Harmansa, Emmanuel Caussinus, Markus Affolter

## Abstract

The *Drosophila* tracheal system consists of an interconnected network of monolayered epithelial tubes that ensures oxygen transport in the larval and adult body. During tracheal dorsal branch (DB) development, individual DBs elongate as a cluster of cells, led by tip cells at the front and trailing cells in the rear. Branch elongation is accompanied by extensive cell intercalation and cell lengthening of the trailing stalk cells. While cell intercalation is governed by Myosin II (MyoII)-dependent forces during tissue elongation in the *Drosophila* embryo leading to germ-band extension, it remained unclear whether MyoII plays a similar active role during tracheal branch elongation and intercalation. Here, we use a nanobody-based approach to selectively knock-down MyoII in tracheal cells. Our data shows that despite the depletion of MyoII function, tip cells migration and stalk cell intercalation (SCI) proceeds at a normal rate. Therefore, our data confirms a model in which DB elongation and SCI in the trachea occurs as a consequence of tip cell migration, which produces the necessary forces for the branching process.

**Summary statement:** Branch elongation during *Drosophila* tracheal development mechanistically resembles MyoII-independent collective cell migration; tensile forces resulting from tip cell migration are reduced by cell elongation and passive stalk cell intercalation.

**Abbreviations:**

DB: Dorsal branch
DC: Dorsal closure
E-Cad: E-Cadherin
GBE: Germ-band extension
MRLC: Myosin regulatory light chain
MyoII: Myosin II
SCI: stalk cell intercalation
Sqh: Spaghetti squash
Sxll: Sex lethal
TC: Tip cell
Tr: Tracheomere

## Introduction

During morphogenesis, a coordinated series of complex events including cell division, cell shape changes and cell rearrangements underlies the formation of functional tissues and organs. Epithelial cell intercalation is a major morphogenetic mechanism acting in polarized tissue elongation, e.g. during *Drosophila* germ-band extension (GBE, Fig. 1A) (Irvine and Wieschaus, 1994), mouse gastrulation (Yen et al., 2009), *C. elegans* intestine (Leung et al., 1999) and *Xenopus* kidney tube development (Lienkamp et al., 2012). During intercalation, controlled cell neighbor-exchange results in tissue extension along one axis and concomitant convergence along the orthogonal axis. Intercalation requires contacts between two adjacent cells to shrink (Fig. 1A’, type I configuration), resulting in a configuration where four or more cells contact each other (type II configuration) (Bertet et al., 2004; Blankenship et al., 2006). Subsequently, the new contact extends (type III configuration) leading to a local extension of the tissue (Bardet et al., 2013; Collinet et al., 2015; Zallen and Wieschaus, 2004).

**Figure 1.**
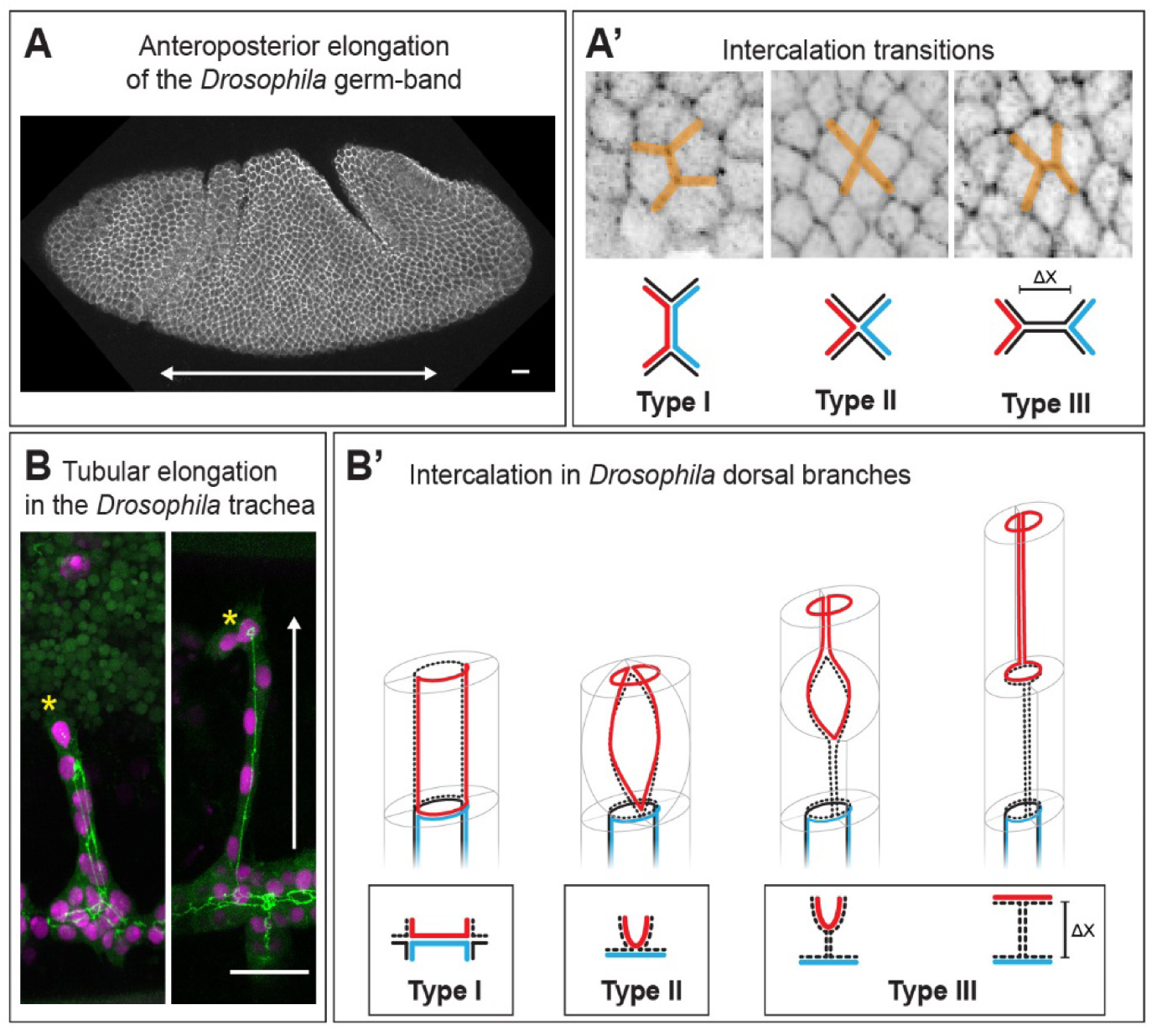
Cell intercalation in flat and tubular epithelia follows similar geometric rules. (A) Anteroposterior elongation of the germ-band (arrow in (A)) in the early *Drosophila* embryo is caused by the intercalation of hexagonal cells. During intercalation and tissue extension junctions remodel in a stereotypic and ordered manner, characterized by three irreversible transitions (A’): Shrinkage of cell contacts between anteriorposterior neighbors (red and blue cell, type I) is followed by new contact formation between dorsal and ventral neighboring cells (black cells, type II). This novel cell-cell contact is expanded (∆x) along the anterior-posterior axis, resulting in net tissue elongation along this axis (type III). (B) Tubular elongation in the *Drosophila* tracheal system. The dorsal branches elongate dorsally (arrow) owing to the migratory behavior of the tip cells (stars) (nuclei in pink, junctions in green). Intercalation in dorsal branches (B’). Pairs of cells remodel their junctions (dashed black and red) during intercalation resulting in a chain-like arrangement of cells after completion of the process. Junction remodeling is polarized and follows a stereotyped pattern corresponding to a type I to type II transition, and junction expansion, corresponding to a type II to type III transition as observed in germ-band extension. (A) Embryo stained for E-Cad, (B) embryo expressing mCherryNLS and α-Catenin GFP driven under the *btl-Gal4* driver.

The *Drosophila* tracheal system presents a paradigm of epithelial remodeling and elongation through cell intercalation in a tubular organ. The primary branches are monolayered epithelial tubes and form in the absence of cell division in two distinct stages. First, tracheal tip cells (TCs) begin to migrate away from the tracheal sac and pull along several tracheal stalk cells into the developing branch, forming a small bud (Samakovlis et al., 1996). In a second phase, the branches elongate and narrow down due to stalk cell intercalation (SCI) and extensive cell lengthening (Ribeiro et al., 2004) (Fig. 1B). SCI of the primary branches follows similar geometrical rules as intercalation in flat epithelia (Lecuit, 2005). Initially, cells in the bud are arranged in a side-to-side configuration and share intercellular junctions with their opposite neighbor but also with cells located distal and proximal along the branch (type I configuration, see Fig. 1B’). Intercalation is initiated by cells reaching around the lumen and forming an autocellular junction (type II configuration), followed by zipping up of the autocellular junction along the proximal-distal axis of the branch (type III configuration). Therefore, the pair of cells initially located side-by-side, rearranges in an end-to-end configuration, resulting in branch elongation (Fig.1B’) (Neumann and Affolter, 2006; Ribeiro et al., 2004).

While the steps of cell and junction rearrangements during intercalation have been described in great detail (see Fig.1), it remains debated whether intercalation *per se* is the driving force leading to branch extension. Several studies in epithelial tissues suggest that intercalation is the direct consequence of increased cortical contractility resulting from the dynamics and the localization of MyoII, thereby generating the major force controlling tissue elongation (Bardet et al., 2013; Bertet et al., 2004; Rauzi et al., 2008; Simoes Sde et al., 2010). However, also external forces acting on tissue boundaries have been implicated in tissue elongation. For instance, extrinsic pulling forces generated by posterior midgut invagination were linked to *Drosophila* GBE (Butler et al., 2009; Collinet et al., 2015; Kong et al., 2016; Lye et al., 2015), also the *Drosophila* wing is shape by extrinsic tensile forces (Etournay et al., 2015; Ray et al., 2015). Therefore, tissue elongation is a consequence of a combination of local and tissue-scale forces. During tracheal dorsal branch (DB) elongation, laser ablation studies have shown that highly motile tip cells create a tensile stress during migration, resulting in branch elongation and SCI (Caussinus et al., 2008). Furthermore, Spaghetti squash-GFP, a GFP fusion of the myosin regulatory light chain (Sqh/MRLC), did not localize to the adherens junctions during SCI. Therefore, and in contrast to elongating epithelial sheets in the fly embryo (see above), cell intercalation appears not to be cause but rather the consequence of epithelial branch elongation in the tracheal system.

Nevertheless, it remains possible that similar to elongating epithelial sheets, a local tensile force at cell boundaries, produced by MyoII activation, plays an additional active role in DB branch elongation. Given the prominent role of MyoII during epithelial morphogenesis, its presence in the tracheal system throughout development and its clear role during earlier tracheal development (Nishimura et al., 2007), we decided to further investigate the function of MyoII during SCI in the tracheal system. To overcome prior limitations due to MyoII maternal contribution and pleiotropic roles in morphogenesis and cytokinesis, we used a nanobody-based approach that can acutely deplete MyoII in a time and tissue specific manner (Caussinus et al., 2012; Pasakarnis et al., 2016). Our results show that in the absence of actomyosin contractibility, tip cell migration and stalk cell intercalation occur normally. Thus our data provides functional evidence supporting a model proposing that primary branch elongation in the trachea is driven by tip cell migration and passive stalk cell intercalation, and demonstrates that the primary tracheal branching process is a consequence of cell migration and thus coordinated by tip cell activity.

## Results

### deGradFP efficiently knocks down MyoII in a time- and tissue-specific manner during embryogenesis

In order to interfere with MyoII function directly at the protein level in a time- and tissue-specific manner, we utilized the deGradFP method. deGradFP allows for the efficient degradation of GFP-fusion proteins and can be used to phenocopy loss-of-function mutations (Blattner et al., 2016; Caussinus et al., 2012; Lee et al., 2016; Nagarkar-Jaiswal et al., 2015; Pasakarnis et al., 2016). Here, we use a null mutant for *sqh* (Jordan and Karess, 1997) rescued by a Sqh-GFP transgene (*sqhAX3*; *sqh-Sqh-GFP*) (Royou et al., 2004). In this genetic background, we expressed deGradFP using the Gal4/UAS system to target Sqh-GFP for degradation in different tissues and analyzed the resulting phenotypes.

In all the experiments shown, the *sqhAX3*; *sqh-Sqh-GFP* line was used as a maternal counterpart in our crossing schemes, allowing us to easily introduce a Gal4 driver and UAS-deGradFP from the paternal side (see Fig.S1 for a detailed description of the crossing schemes). Since *sqh* is on the X-chromosome, all male progeny from a cross were hemizygous for *sqhAX3* and hence the *sqh-Sqh-GFP* transgene on the second chromosome provided the only source of Sqh protein. In contrast, females expressed both non-tagged and GFP-tagged Sqh. In order to distinguish male embryos from female embryos, we used different approaches depending on the experimental condition. In fixed embryos, we used a monoclonal antibody against Sex lethal (Sxl), which recognizes all somatic cell nuclei in females from nuclear cycle 12 (Bopp et al., 1991). For live imaging analyses, we used a *vestigial* red fluorescent reporter (5XQE-DsRed) on the X chromosome that has a segmental pattern of expression in the epidermis from stage 11 and continues to be expressed in the embryo and in larval stages in diverse tissues (Zecca and Struhl, 2007). These two methods, in combination with fluorescent reporters and the obvious signs of Sqh-GFP degradation (see below), allowed us to unambiguously discriminate all possible genotypes.

Initially, we validated the efficiency of deGradFP-mediated knock-down of Sqh-GFP in the lateral epidermis, due to its imaging accessibility and comprehensive characterization. During dorsal closure (DC), epidermal cells elongate in dorso-ventral direction to close a gap that exists in the dorsal epidermis. Along the leading edge of the closing epidermis, a supra-cellular actomyosin cable forms (Fig.2A) (Franke et al., 2005; Kiehart et al., 2000; Pasakarnis et al., 2016) and leading cells project filopodia and lamellipodia dorsally (Fig.2A, right) (Eltsov et al., 2015). To perturb Sqh function in the embryonic epidermis, we used *engrailed*-Gal4 (*en*-Gal4) (Tabata et al., 1992) to restrict expression of deGradFP to the posterior compartment of each segment in *sqhAX3*; *sqh-Sqh-GFP* embryos.

**Figure 2.**
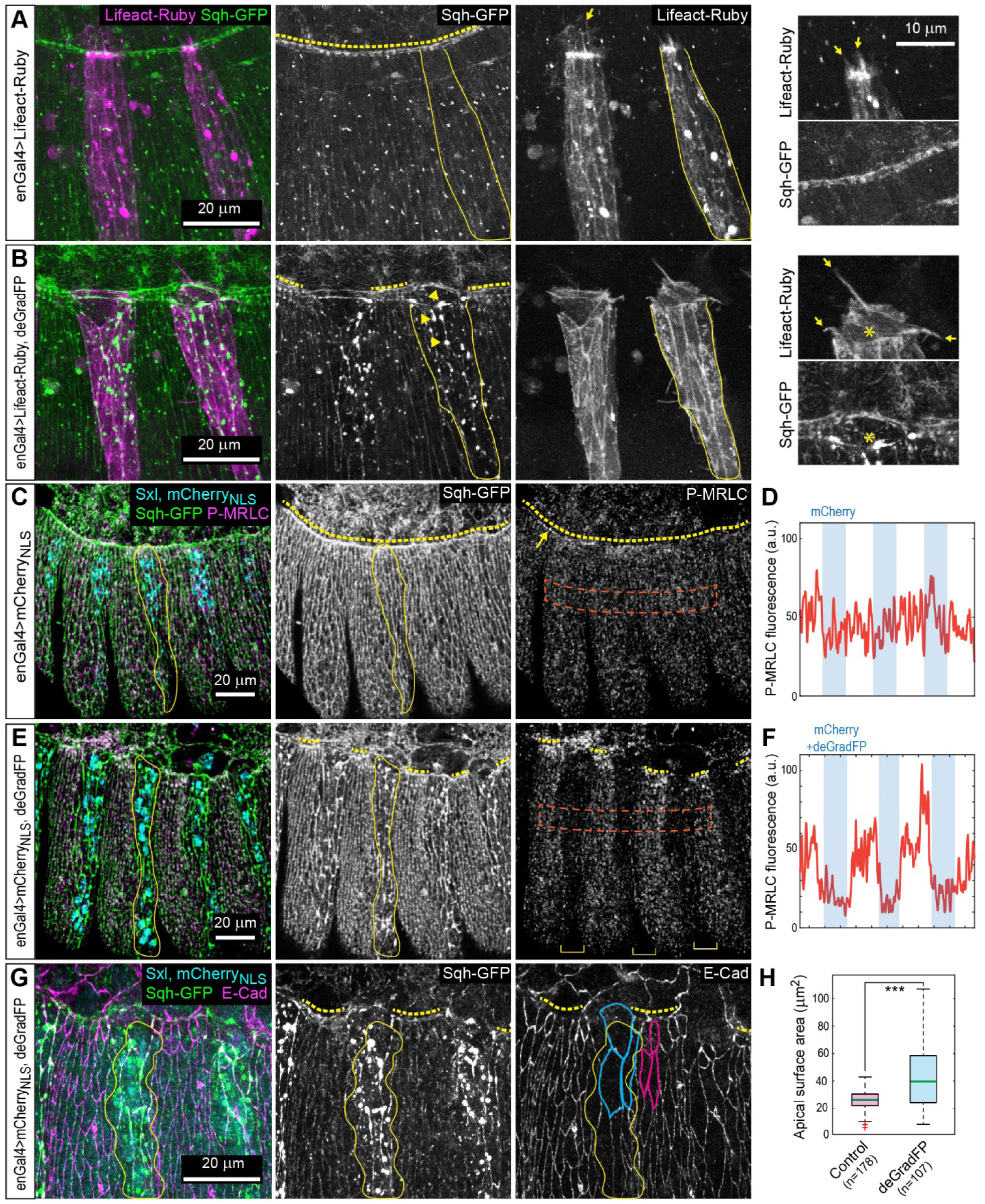
deGradFP-mediated knock-down of Sqh-GFP in the embryonic epidermis. All panels show lateral views of stage 14 male *sqhAX3*; *sqh-Sqh-GFP* embryos additionally expressing the indicated transgenes in the engrailed (*en::Gal4*) stripe pattern. Representative *en* stripes are highlighted by continuous yellow lines. (A, B) Live imaging revealed that in control embryos (A) Sqh-GFP localizes in small punctuae at the cell cortex and forms a continuous actomyosin cable (yellow dashed line). In contrast, in deGradFP expressing embryos Sqh-GFP coalesces prominently at the cortex forming large spots (arrowheads in (B) middle) and the dorsal actomyosin cable in lost the *en* stripes. Also, deGradFP expressing embryos formed more and longer filopodia at the leading epidermal edge visualized by Lifeact-Ruby (compare arrows in (A, B) right and magnification to the right). (C-E) Control embryos stained for phosphorylated MRLC (P-MRLC) show uniform phosphomyosin distribution in all segments and enrichment at the actomyosin cable (arrow in (C). While in deGradFP expressing embryos the P-MRLC signal is drastically reduced in en stripes (see yellow brackets in (E) and (F). P-MRLC fluorescence of the areas marked by a red dashed line (in C, E) are plotted in (D, F), respectively. (G) deGradFP expressing embryo stained for E-Cadherin (E-Cad) shows increased apical surface area in the *en* stripe (blue outlines) compared to cells outside of *en* stripes (pink outlines). (H) Quantification of apical cell surface area of cells inside (blue) and outside (pink) of the *en* stripe. The green lines mark the median; whiskers correspond to minimum and maximum data points. Statistical significance was assessed using a two-sided Students *t-test* (*** *p*<0.001), outliers are indicated by a red cross.

deGradFP mediated Sqh-GFP knock-down resulted in an interruption of the actomyosin cable. However, deGradFP mediated knock-down did not result in the total disappearance of Sqh-GFP, instead Shq-GFP remained in what appears to be inclusion bodies (Fig.2B, arrowheads). This observation was similar to what was previously reported in the epidermis (Pasakarnis et al., 2016) and the wing imaginal discs (Caussinus et al., 2012). Live imaging of the F-actin reporter Lifeact-Ruby revealed that Sqh-GFP knock-down males formed thicker and longer filipodia at the leading edge than the control embryos (compare Fig.2A and B, yellow arrows). However, F-actin was not enriched at the bright Sqh-GFP “inclusion bodies” (Fig.2B, asterisk, see also Pasakarnis et al., 2016).

To ensure that Sqh function was indeed lost under these conditions, we used an antibody specifically recognizing the phosphorylated and active form of Sqh (P-MRLC) (Ikebe and Hartshorne, 1985; Jordan and Karess, 1997; Karess et al., 1991). In stage 14 male control embryos, P-MRLC showed apical punctate localization and enrichment at the actomyosin cable (Fig.2C, arrow and Fig.2D). In knock-down embryos, P-MRLC levels were strongly reduced in all deGradFP expressing cells, even at the leading edge (Fig.2E, F). Furthermore, it was suggested previously that a loss of MyoII function results in cortical relaxation (Mason et al., 2013; Royou et al., 2002; Rozbicki et al., 2015). We therefore investigated whether knock-down of Sqh-GFP resulted in aberrant cell morphology. Indeed, staining for the junctional protein E-Cadherin (E-Cad) revealed that Sqh-GFP knock-down resulted in a significant increase in apical cell surface area (Fig.2G, H and Fig.S2).

By stage 16, Sqh-GFP knock-down cells managed to contact cells in the contralateral stripes. We observed that in the posterior half of the embryo, cells in the deGradFP expressing stripes moved forward, displacing the non-deGradFP expressing cells and excluding them completely from the leading edge while sealing aberrantly with other deGradFP expressing cells and never with non-deGradFP expressing cells (Fig.S4C). These results are similar to the ones obtained by the striped expression of a dominant negative version of Rho1, a positive upstream regulator of MyoII (Jacinto et al., 2002) and two more recent studies using either loss-of-function mutants or deGradFP to disrupt the leading edge actomyosin cable (Ducuing and Vincent, 2016; Pasakarnis et al., 2016). Finally, knock-down of Sqh-GFP in stripes resulted in embryonic lethality in male embryos, while female embryos, carrying a wild type *sqh* copy, gave rise to viable progeny (Fig.S3 and Fig.S4B).

In summary, these results consistently show that deGradFP-mediated knock-down of Sqh-GFP in the rescue background inactivates MyoII and produces phenotypes consistent with a loss of MyoII, namely abnormal epidermal packing, disruption of the actomyosin cable and aberrant epidermal leading edge behavior, without disrupting other actin based structures such as filopodia. Furthermore, we have tested a number of other means to inactivate MyoII function but did not find another method that resulted in dorsal open phenotypes as observed when knocking down Sqh-GFP in amnioserosa cells using deGradFP (Table S1). Therefore, we conclude that deGradFP represents the best available tool to address the role of MyoII during cell intercalation in tracheal branch formation.

### Tracheal specific knock-down of Sqh/MRLC does not perturb primary branch elongation

To determine whether MyoII function is necessary for tracheal system development, we expressed deGradFP under control of the trachea specific *btl-Gal4* driver in *sqh* mutant background. *btl-Gal4* is expressed in tracheal cells from stage 11 onward, after the invagination of the tracheal placode. Low magnification time lapse imaging of knock-down embryos revealed that Sqh-GFP showed a dotted appearance at each tracheomere, indicating deGradFP activity and efficient inactivation of Sqh-GFP (Movie 1). E-Cad staining in fixed embryos showed that the overall development and the morphology of the tracheal system remained normal upon expression of deGradFP; all major tracheal branches formed and fused with their corresponding partners, similar to what is observed in control embryos (Fig.3A, B). Additionally, the deposition and clearance of the tracheal chitin cable and the subsequent gas filling of the tracheal tubes proceeded normally in knock-down embryos (Fig.S5 and Movie 2). Using the markers described above, we verified that the full knock-down embryos were male and died as first instar larvae while female embryos where viable, further validating the absence of a dominant negative effect of degrading Sqh-GFP in the presence of a *sqh* wild type gene.

**Figure 3.**
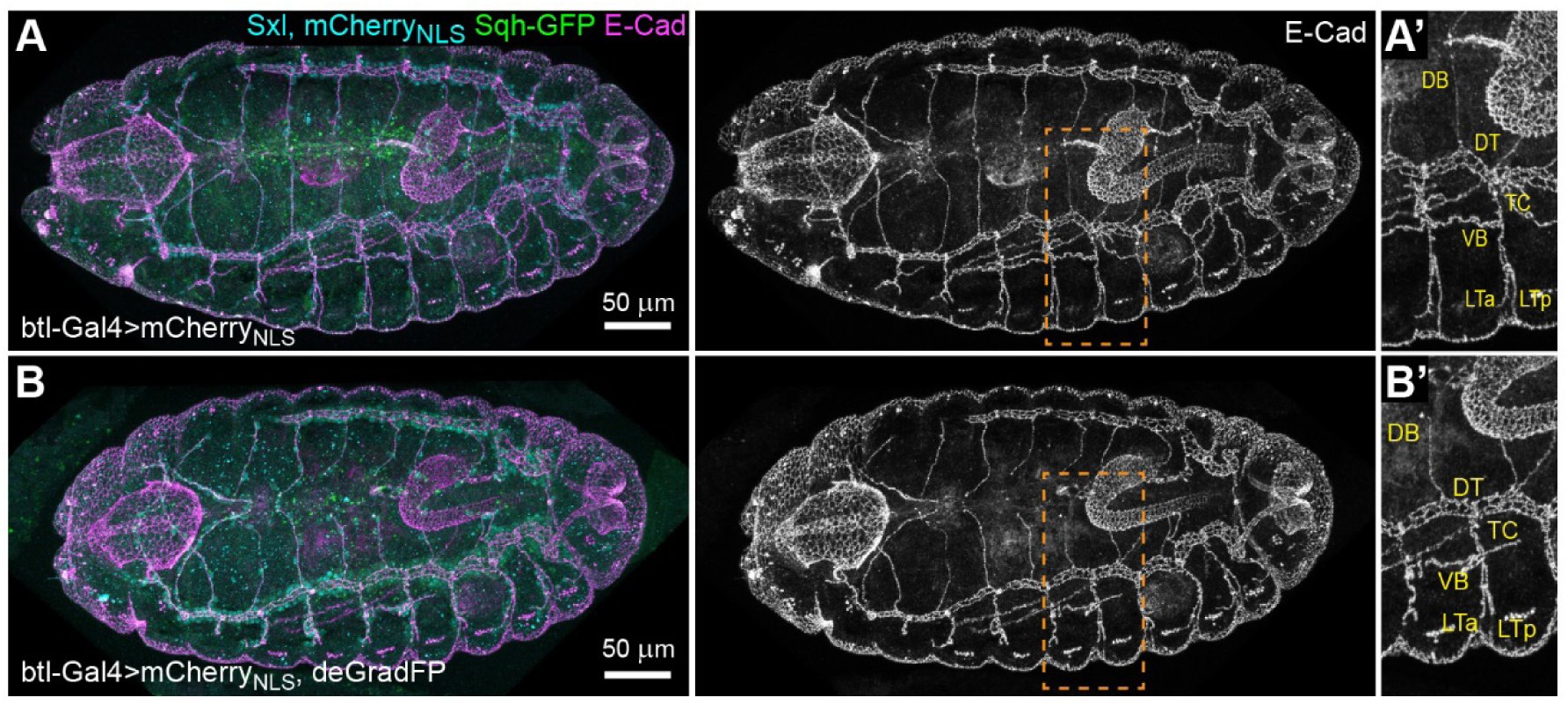
Sqh-GFP knock-down in the trachea system results in normal branch architecture. All panels show dorsal views of stage 16 male *sqhAX3*; *sqh-Sqh-GFP* embryos expressing mCherry_NLS_ (A, control) or mCherryNLS together with deGradFP (B) in the tracheal system (*btl-Gal4*). The trachea system architecture is visualized by staining for E-Cad. In male control (A) and tracheal Sqh knock-down (B) embryos all main tracheal branches form, elongate and fuse to develop a normal tracheal system. In the magnification (A’, B’) the tracheomere 5 (Tr5) branches are labelled: dorsal branch (DB), dorsal trunk (DT), transverse connective (TC), visceral branch (VB) and lateral trunk branches (LTa and LTp).

To gain a more detailed view of possible consequences of the absence of MyoII activity during cell intercalation, we characterized the dynamics of DB elongation upon Sqh-GFP knock-down in tracheal cells from stage 11 onward (*btl-Gal4*). Time lapse movies of male *sqhAX3*; *sqh-Sqh-GFP* embryos expressing either mCherryNLS alone (Control) or mCherryNLS and deGradFP (deGradFP) allowed us to investigate the dynamics of DB elongation (Fig.4A-D and Movie 3). In both conditions, 5-6 DB cells were present in an initial side by side configuration (Fig.4A, B 0min., also see Fig.S6). In the following elongation phase, tip cells migrated dorsally (Fig.4A, B 35 min.) while the stalk cells intercalated, i.e. the cells rearranged to an end-to-end configuration (Fig.4A, B 125 min.). The intercalation process and the dynamics we observed during DB elongation in knock-down embryos were indistinguishable from control embryos (Fig.4C-D). Furthermore, time lapse movies of male *sqhAX3*; *sqh-Sqh-GFP* embryos expressing either Lifeact-Ruby alone (Control) or Lifeact-Ruby and deGradFP (deGradFP) showed that knock-down DBs tip cells formed filopodia (Fig.4E, F and Movie 4) similar to wild type embryos (Lebreton and Casanova, 2014; Ribeiro et al., 2004). In later stages, terminal cells formed and the fusion cells contacted the contralateral DB in the Sqh-GFP knock-down embryos, comparable to control embryos (Fig.4E, F 160 min).

**Figure 4.**
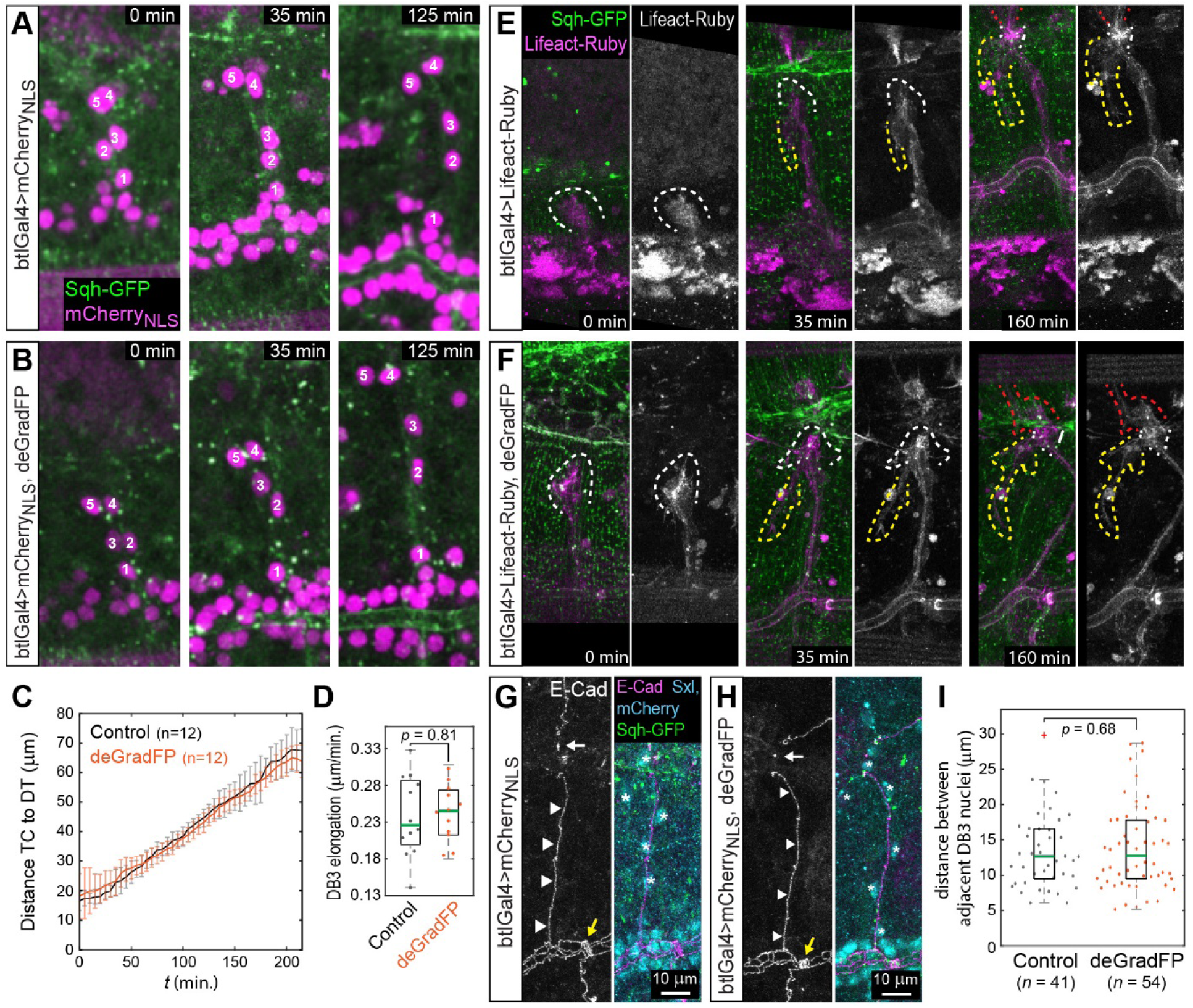
Dorsal branch elongation and SCI does not require MyoII activity. All panels show male *sqhAX3*; *sqh-Sqh-GFP* embryos expressing the indicated transgenes in the tracheal system (*btl-Gal4*). (A, B) Stills from time lapse movies show that in control (A) and Sqh-GFP knock-down embryos (B) DBs elongate (individual cells are numbered). (C) Distance between the tip cell (TC) and the dorsal trunk (DT) during Tr3 DB elongation. Tr3 DB elongation starts to plateau after 200 min. at approximately 65μm in both conditions. Error bars indicate the standard deviation. (D) Rate of DB elongation. The median is marked by a green line. (E, F) In both, control (E) and Sqh knock-down embryos (F), tip cells extend filopodia (white dashed lines 0min. and 35min.), form a terminal cell with long cytoplasmic extensions (yellow dashed lines in 35min. and 160min.) and form a fusion cell that eventually contacts the contralateral branch (white and red dashed lines 160min). (G, H) Control (G) and Sqh-GFP knock-down (H) stage 16 embryos showing the Tr3 DBs stained for E-Cad. In both cases cells within the branch have an end to end configuration (asterisks over nuclei), form autocellular junctions (arrowheads) and deposit de novo junctional material in contact with the contralateral branch (white arrows). Also highlighted are the fused Tr3 and Tr4 in the DT (yellow arrows). (I) Distance between individual nuclei in the Tr3 DBs of stage 16 embryos. Median (green line), whiskers correspond to minimum and maximum data points. Statistical significance was assessed using a two-sided Students t-test, outliers are indicated by a red cross.

Finally, E-Cad staining on control and knock-down embryos revealed that in stage 16 knock-down embryos (in which nuclei had an end-to-end arrangement), intercellular junctions had remodeled to give rise to autocellular junctions, again as seen in control embryos (Fig.4G, H). In this final stage, the fusion cells established de novo contacts with the contralateral branch, as visualized by a dot of E-Cad between the two fusion cells in both conditions (Fig.4G, H). Also, the spacing between DB nuclei in stage 16 embryos did not significantly differ between the two conditions (Fig.4I). These results show that MyoII activity is not required during dorsal branch migration, elongation and fusion, and more importantly, the junctional remodeling in SCI proceeds normally in the absence of functional MyoII.

## Discussion

A long standing question in the field was whether MyoII activity is required for DB elongation and SCI. Here we present data showing that MyoII function is dispensable for branch elongation and concomitant SCI during *Drosophila* tracheal development. Acute depletion of Sqh-GFP, the regulatory light chain of MyoII, specifically in the tracheal system resulted in a normal architecture of the tracheal system and the dynamics of DB elongation and SCI were unaffected. The experiments and results presented here are in line with our previous observations that the pulling forces provided by the tip cells provide enough mechanical force for SCI, and fully support a scenario in which stalk cell intercalation is a cell non-autonomous process brought about by tip cell migration (Caussinus et al., 2008).

Critical to our approach and different from previous ones was the use of deGradFP to acutely deplete Sqh-GFP protein in order to block MyoII function. Previous approaches mainly relied on hypomorphic mutants and overexpression of dominant-negative and inhibitory proteins in order to interfere with MyoII function. However, these approaches hold drawbacks complicating the interpretation of the experimental outcomes. Mutants used to study MyoII function during late embryogenesis must not interfere with maternally contributed mRNA and protein (Franke et al., 2010) in order to allow normal early embryonic development. Hence, due to protein stability, MyoII function might not be completely lost in such a background. Furthermore, mutations often affect multiple cellular processes and therefore are prone to generate indirect effects or lead to adaptation. To overcome these drawbacks, time- and tissue-specific expression of dominant negative forms of MyoII or upstream regulators have often been used (Fischer et al., 2014; Franke et al., 2010; Saias et al., 2015). However, these tools seemed to be less efficient in their depleting competence and gave rise to much milder phenotypes than the ones observed with deGradFP (see also Pasakarnis et al., 2016 and Table S1). Therefore, deGradFP is the most effective tool available to deplete Sqh-GFP and interfere with MyoII function in a time- and tissue-specific manner.

The results we obtain by acute depletion of Sqh-GFP in the trachea system provide two unexpected findings. First, cellular rearrangements during SCI occur normally in the absence of MyoII activity, and second, actomyosin contractile forces are not required in tracheal cells for TC migration and concomitant DB elongation.

The forces that fuel epithelial cell intercalation and tissue elongation have been intensely studied in several organisms. During *Drosophila* germ-band extension, local forces arising from spatiotemporal dynamics in MyoII levels are required for junctional shrinkage (Bertet et al., 2004; Blankenship et al., 2006; Fernandez-Gonzalez et al., 2009; Levayer and Lecuit, 2013; Rauzi et al., 2008) and subsequent extension (Bardet et al., 2013; Collinet et al., 2015) and act together with global, tissue scale forces (Butler et al., 2009; Collinet et al., 2015; Etournay et al., 2015; Ray et al., 2015) to drive tissue elongation. Therefore, cell intercalation is a direct consequence of local and tissue scale forces and a major cause of tissue elongation in the *Drosophila* germ-band. MyoII activity was also shown to be required for intercalation during chicken primitive streak formation (Rozbicki et al., 2015), and mouse renal tube elongation (Lienkamp et al., 2012). Therefore, most intercalation processes mechanistically closely resemble GBE in the *Drosophila* embryo and use locally produced forces to drive junction and cell-neighbor remodeling. In contrast to the control of intercalation by local force development, external constrains acting on tissue boundaries also control and/or drive intercalation and tissue remodeling. This is very likely the case during *Drosophila* pupal wing extension, where anchorage of wing blade cells to the pupal cuticle and synchronous contraction of the hinge create a tissue scale force pattern that drives cellular rearrangements via intercalation and cell division (Etournay et al., 2015; Ray et al., 2015). Our results suggest that despite the resemblance of tracheal SCI to intercalation in the embryonic epidermis (see Fig. 1), the molecular mechanisms underlying force generation in these two systems are fundamentally different.

Pulling forces during tracheal branch elongation arise due to TC migration, which result in an extrinsic traction force that creates tension in the trailing stalk cells (Caussinus et al., 2008). Cell elongation and rearrangements associated with oriented cell division were shown to result in stress dissipation (Affolter et al., 2009; Campinho et al., 2013; Guillot and Lecuit, 2013; Wyatt et al., 2015). Since tracheal branch elongation occurs in the absence of cell division, extensive cell elongation and cell intercalation presumably provide the only mechanisms for tension relaxation in this context. This is in line with the earlier proposal that cell shape changes and SCI are passive, non-cell autonomous processes induced by the tension created in stalk cells by TC migration (also see Affolter and Caussinus, 2008; Affolter et al., 2009). Therefore, while during GBE locally produced, MyoII-dependent, forces drive intercalation and tissue extension, our results show that SCI in the trachea is MyoII-independent and that tracheal branch intercalation is, similar to pupal wing extension (Etournay et al., 2015; Ray et al., 2015), a passive process driven by global tissue scale pulling forces.

Despite the loss of actomyosin activity in all tracheal cells, including the TCs, we found that branch elongation dynamics are unchanged. Therefore, TC migration does not rely on actomyosin contractibility, posing the question which molecular players might be involved in TC force generation. Interestingly, collective movement of cells depends on actin-based filopodia and lamellipodia (Mayor and Etienne-Manneville, 2016) and was shown in several cases to be independent of MyoII activity (Matsubayashi et al., 2011; Serra-Picamal et al., 2012). Furthermore, several studies showed that a down-regulation of MyoII is required for effective collective cell migration (Hidalgo-Carcedo et al., 2011; Omelchenko and Hall, 2012; Yamada and Nelson, 2007). It seems that MyoII-independent TC migration might be mainly actin polymerization-based and the branching process might therefore be more similar to collective cell migration than to classical epithelial intercalation in flat tissues such as the embryonic epidermis.

Future studies will need to investigate the detailed molecular basis of force generation during trachea TC migration. The extent to which TC migration and DB elongation depends on actin polymerization and the molecules participating in this process might be investigated by directly modulating actin polymerization regulators using deGradFP.

## Materials and Methods

### *Drosophila* stocks

The following stocks were used: *btl-Gal4* (Shiga et al., 1996), UAS-mCherry_NLS_ (Caussinus et al., 2008), UAS-LifeAct-Ruby (Hatan et al., 2011), UAS-deGradFP (Caussinus et al., 2012), 5XQEDSRed (Zecca and Struhl, 2007), *sqhAX3*; *sqh-Sqh-GFP* (Royou et al., 2004), dominant negative version of zip (UAS-GFP-DN-zip, Franke et al., 2005), *en-Gal4*, *amnioserosa-Gal4* ({PGawB}332.3), UAS-Dicer2, UAS-shRNA-sqh (TRiP.HMS00437, TRiP.HMS00830 and TRiP.GL00663), UAS-shRNA-zip (TRiP.HMS01618 and TRiP.GL00623), dominant negative version of Rok (UAS-rok.CAT-KG2B1 and UAS-rok.CAT-KG3 (Winter et al., 2001)), dominant negative version of Rho1 (UAS-Rho1.N19, Strutt et al., 1997) (Bloomington Stock Center).

### Immunohistochemistry and antibodies

The following antibodies were used: mouse anti-Sxl-m18 (1:100; DSHB), rabbit anti Phospho-Myosin Light Chain 2 (Ser19) (1:50; #3671 Cell Signaling Technology), rabbit anti-Verm (1:300, gift from S. Luschnig), rat anti E-Cad DCAD2 (1:100; DSHB). Secondary antibodies were conjugated with Alexa 488, Alexa 568, Alexa 633 (Molecular probes) or Cy5 (Jackson ImmunoResearch). Embryos were collected overnight and fixed in 4% formaldehyde in PBS-heptane for 20 min or 10 min (for anti E-Cad) and devitellinized by shaking in methanol-heptane. After extensive washing in methanol and PBT, embryos were blocked in PBT containing 2% normal goat serum and incubated in primary antibody solution overnight at 4°C. The next day, embryos were extensively washed with PBT and incubated in primary antibody solution for 2 hours at room temperature. Subsequently embryos were washed in PBT again and mounted in Vectashield (H-1000, Vector Laboratories).

### Light microscopy

Imaging was carried out using a Leica TCS SP5 confocal microscope with x20 dry, x40W x63W and x63glicerol objectives. For live imaging embryos were collected overnight, dechorionated in 4% bleach, and mounted in 400-5 mineral oil (Sigma Diagnostic, St Louis, MO) between a glass coverslip and a gas-permeable plastic foil (bioFOLIE 25, In Vitro System and Services, Gottingen, Germany). Imaging was done at 10 min. intervals for movie 1 and 2, at 5 min. intervals for movie 3 and at 2 min. intervals for movie 4. Images were processed using ImageJ (v1.42; NIH) and Imaris (v7.3.0; Bitplane). Time-lapse movies were processed using a custom made plugin in ImageJ to correct for drift in the xy plane.

### Quantifications and statistics

For the P-MRLC plots in Fig.2D, F we measured the fluorescent intensities in the regions of interest indicated in Fig.2C, E using the Plot Profile function in ImageJ (NIH). Apical cell surface area (quantifications shown in Fig.2H and Fig.S2B) was measured in ImageJ from maximum projections of pre-DC stage 15 embryos stained for E-Cad. We excluded cells from the quantification that we could not clearly assign to either the En positive (mCherryNLS) or the En negative stripes. To quantify the dynamics of branch elongation (Fig.4C, D), we measured the direct (minimal) distance between the dorsal trunk and the tip cells of the Tr3 DB in maximum projections of time-lapse movies using ImageJ. The plot in Fig.4C shows the arithmetic means and the error bars show the standard deviation. For the quantifications in Fig.4I, live embryos were collected and staged using completion of dorsal closure as a reference to obtain stage 16 embryos. Live embryos were mounted dorsolaterally as previously described and only embryos in which dorsal branches nuclei appeared in the same plane were imaged at 1 μm optical section intervals. Z maximum projections of the acquired images were used to measure the distances between nuclei in dorsal branch 3 (which migrates the longest distance in wt) using imageJ. n-numbers are indicated either directly in the figures or in the corresponding legend. In the boxplots (Fig.2H, Fig.4D, I and Fig.S2B) center values (green bar) correspond to the median and whiskers mark maximum and minimum data points. Sample number was chosen large enough to allow statistical significance being assessed using a two-sided Student’s *t*-test with unequal variance.

## Acknowledgements

We thank the Bloomington Stock Center, the Developmental Studies Hybridoma Bank, Gary Struhl and Stefan Luschnig for providing fly stocks and antibodies. The Biozentrum Imaging Core Facility for maintenance of microscopes and support. We thank MM. Baer, A. Lenard, O. Kanca, F. Hamaratoglu, AS. Denes and M. Müller for helpful discussions and technical assistance. M. Brauchle for advice and comments on the manuscript.

## Author contributions

A.O., E.C. and M.A. designed the study, A.O. and E.C. performed the experiments, A.O., E.C. and S.H. analyzed and quantified the data, all authors contributed to the writing of the paper.

## References

Affolter, M., Caussinus, E. (2008). Tracheal branching morphogenesis in Drosophila: new insights into cell behaviour and organ architecture. Development 135, 2055–2064.

Affolter, M., Zeller, R., Caussinus, E. (2009). Tissue remodelling through branching morphogenesis. Nature reviews. Molecular cell biology 10, 831–842.

Bardet, P.L., Guirao, B., Paoletti, C., Serman, F., Leopold, V., Bosveld, F., Goya, Y., Mirouse, V., Graner, F., Bellaiche, Y. (2013). PTEN Controls Junction Lengthening and Stability during Cell Rearrangement in Epithelial Tissue (vol 25, pg 534, 2013). Developmental cell 26, 674–674.

Bertet, C., Sulak, L., Lecuit, T. (2004). Myosin-dependent junction remodelling controls planar cell intercalation and axis elongation. Nature 429, 667–671.

Blankenship, J.T., Backovic, S.T., Sanny, J.S.P., Weitz, O., Zallen, J.A. (2006). Multicellular rosette formation links planar cell polarity to tissue morphogenesis. Developmental cell 11, 459–470.

Blattner, A.C., Chaurasia, S., McKee, B.D., Lehner, C.F. (2016). Separase Is Required for Homolog and Sister Disjunction during Drosophila melanogaster Male Meiosis, but Not for Biorientation of Sister Centromeres. PLoS genetics 12, e1005996.

Bopp, D., Bell, L.R., Cline, T.W., Schedl, P. (1991). Developmental distribution of female-specific Sex-lethal proteins in Drosophila melanogaster. Genes & development 5, 403–415.

Butler, L.C., Blanchard, G.B., Kabla, A.J., Lawrence, N.J., Welchman, D.P., Mahadevan, L., Adams, R.J., Sanson, B. (2009). Cell shape changes indicate a role for extrinsic tensile forces in Drosophila germ-band extension. Nature cell biology 11, 859–864.

Campinho, P., Behrndt, M., Ranft, J., Risler, T., Minc, N., Heisenberg, C.P. (2013). Tension-oriented cell divisions limit anisotropic tissue tension in epithelial spreading during zebrafish epiboly. Nature cell biology 15, 1405–1414.

Caussinus, E., Colombelli, J., Affolter, M. (2008). Tip-cell migration controls stalk-cell intercalation during Drosophila tracheal tube elongation. Current biology: CB 18, 1727–1734.

Caussinus, E., Kanca, O., Affolter, M. (2012). Fluorescent fusion protein knockout mediated by anti-GFP nanobody. Nature structural & molecular biology 19, 117–121.

Collinet, C., Rauzi, M., Lenne, P.F., Lecuit, T. (2015). Local and tissue-scale forces drive oriented junction growth during tissue extension. Nature cell biology 17, 1247-+.

Ducuing, A., Vincent, S. (2016). The actin cable is dispensable in directing dorsal closure dynamics but neutralizes mechanical stress to prevent scarring in the Drosophila embryo. Nature cell biology 18, 1149–1160.

Eltsov, M., Dube, N., Yu, Z., Pasakarnis, L., Haselmann-Weiss, U., Brunner, D., Frangakis, A.S. (2015). Quantitative analysis of cytoskeletal reorganization during epithelial tissue sealing by large-volume electron tomography. Nature cell biology 17, 605–614.

Etournay, R., Popovic, M., Merkel, M., Nandi, A., Blasse, C., Aigouy, B., Brandl, H., Myers, G., Salbreux, G., Julicher, F., et al. (2015). Interplay of cell dynamics and epithelial tension during morphogenesis of the Drosophila pupal wing. eLife 4, e07090.

Fernandez-Gonzalez, R., Simoes Sde, M., Roper, J.C., Eaton, S., Zallen, J.A. (2009). Myosin II dynamics are regulated by tension in intercalating cells. Developmental cell 17, 736–743.

Fischer, S.C., Blanchard, G.B., Duque, J., Adams, R.J., Arias, A.M., Guest, S.D., Gorfinkiel, N. (2014). Contractile and mechanical properties of epithelia with perturbed actomyosin dynamics. PloS one 9, e95695.

Franke, J.D., Montague, R.A., Kiehart, D.P. (2005). Nonmuscle myosin II generates forces that transmit tension and drive contraction in multiple tissues during dorsal closure. Current biology: CB 15, 2208–2221.

Franke, J.D., Montague, R.A., Kiehart, D.P. (2010). Nonmuscle myosin II is required for cell proliferation, cell sheet adhesion and wing hair morphology during wing morphogenesis. Developmental biology 345, 117–132.

Guillot, C., Lecuit, T. (2013). Mechanics of epithelial tissue homeostasis and morphogenesis. Science 340, 1185–1189.

Hatan, M., Shinder, V., Israeli, D., Schnorrer, F., Volk, T. (2011). The Drosophila blood brain barrier is maintained by GPCR-dependent dynamic actin structures. The Journal of cell biology 192, 307–319.

Hidalgo-Carcedo, C., Hooper, S., Chaudhry, S.I., Williamson, P., Harrington, K., Leitinger, B., Sahai, E. (2011). Collective cell migration requires suppression of actomyosin at cell-cell contacts mediated by DDR1 and the cell polarity regulators Par3 and Par6. Nature cell biology 13, 49–58.

Ikebe, M., Hartshorne, D.J. (1985). Phosphorylation of smooth muscle myosin at two distinct sites by myosin light chain kinase. The Journal of biological chemistry 260, 10027–10031.

Irvine, K.D., Wieschaus, E. (1994). Cell Intercalation during Drosophila Germband Extension and Its Regulation by Pair-Rule Segmentation Genes. Development 120, 827–841.

Jacinto, A., Wood, W., Woolner, S., Hiley, C., Turner, L., Wilson, C., Martinez-Arias, A., Martin, P. (2002). Dynamic analysis of actin cable function during Drosophila dorsal closure. Current biology: CB 12, 1245–1250.

Jordan, P., Karess, R. (1997). Myosin light chain-activating phosphorylation sites are required for oogenesis in Drosophila. The Journal of cell biology 139, 1805–1819.

Karess, R.E., Chang, X.J., Edwards, K.A., Kulkarni, S., Aguilera, I., Kiehart, D.P. (1991). The regulatory light chain of nonmuscle myosin is encoded by spaghetti-squash, a gene required for cytokinesis in Drosophila. Cell 65, 1177–1189.

Kiehart, D.P., Galbraith, C.G., Edwards, K.A., Rickoll, W.L., Montague, R.A. (2000). Multiple forces contribute to cell sheet morphogenesis for dorsal closure in Drosophila. The Journal of cell biology 149, 471–490.

Kong, D., Wolf, F., Grosshans, J. (2016). Forces directing germ-band extension in Drosophila embryos. Mechanisms of development.

Lebreton, G., Casanova, J. (2014). Specification of leading and trailing cell features during collective migration in the Drosophila trachea. Journal of cell science 127, 465–474.

Lecuit, T. (2005). Adhesion remodeling underlying tissue morphogenesis. Trends in cell biology 15, 34–42.

Lee, K.H., Zhang, P., Kim, H.J., Mitrea, D.M., Sarkar, M., Freibaum, B.D., Cika, J., Coughlin, M., Messing, J., Molliex, A., et al. (2016). C9orf72 Dipeptide Repeats Impair the Assembly, Dynamics, and Function of Membrane-Less Organelles. Cell 167, 774–788 e717.

Leung, B., Hermann, G.J., Priess, J.R. (1999). Organogenesis of the Caenorhabditis elegans intestine. Developmental biology 216, 114–134.

Levayer, R., Lecuit, T. (2013). Oscillation and polarity of E-cadherin asymmetries control actomyosin flow patterns during morphogenesis. Developmental cell 26, 162–175.

Lienkamp, S.S., Liu, K., Karner, C.M., Carroll, T.J., Ronneberger, O., Wallingford, J.B., Walz, G. (2012). Vertebrate kidney tubules elongate using a planar cell polarity-dependent, rosette-based mechanism of convergent extension. Nature genetics 44, 1382–1387.

Lye, C.M., Blanchard, G.B., Naylor, H.W., Muresan, L., Huisken, J., Adams, R.J., Sanson, B. (2015). Mechanical Coupling between Endoderm Invagination and Axis Extension in Drosophila. PLoS biology 13, e1002292.

Mason, F.M., Tworoger, M., Martin, A.C. (2013). Apical domain polarization localizes actin-myosin activity to drive ratchet-like apical constriction. Nature cell biology 15, 926–936.

Matsubayashi, Y., Razzell, W., Martin, P. (2011). 'White wave' analysis of epithelial scratch wound healing reveals how cells mobilise back from the leading edge in a myosin-II-dependent fashion. Journal of cell science 124, 1017–1021.

Mayor, R., Etienne-Manneville, S. (2016). The front and rear of collective cell migration. Nat Rev Mol Cell Bio 17, 97–109.

Nagarkar-Jaiswal, S., Lee, P.T., Campbell, M.E., Chen, K., Anguiano-Zarate, S., Gutierrez, M.C., Busby, T., Lin, W.W., He, Y., Schulze, K.L., et al. (2015). A library of MiMICs allows tagging of genes and reversible, spatial and temporal knockdown of proteins in Drosophila. eLife 4.

Neumann, M., Affolter, M. (2006). Remodelling epithelial tubes through cell rearrangements: from cells to molecules. Embo Rep 7, 36–40.

Nishimura, M., Inoue, Y., Hayashi, S. (2007). A wave of EGFR signaling determines cell alignment and intercalation in the Drosophila tracheal placode. Development 134, 4273–4282.

Omelchenko, T., Hall, A. (2012). Myosin-IXA regulates collective epithelial cell migration by targeting RhoGAP activity to cell-cell junctions. Current biology: CB 22, 278–288.

Pasakarnis, L., Frei, E., Caussinus, E., Affolter, M., Brunner, D. (2016). Amnioserosa cell constriction but not epidermal actin cable tension autonomously drives dorsal closure. Nature cell biology.

Rauzi, M., Verant, P., Lecuit, T., Lenne, P.F. (2008). Nature and anisotropy of cortical forces orienting Drosophila tissue morphogenesis. Nature cell biology 10, 1401–1410.

Ray, R.P., Matamoro-Vidal, A., Ribeiro, P.S., Tapon, N., Houle, D., Salazar-Ciudad, I., Thompson, B.J. (2015). Patterned Anchorage to the Apical Extracellular Matrix Defines Tissue Shape in the Developing Appendages of Drosophila. Developmental cell 34, 310–322.

Ribeiro, C., Neumann, M., Affolter, M. (2004). Genetic control of cell intercalation during tracheal morphogenesis in Drosophila. Current Biology 14, 2197–2207.

Royou, A., Field, C., Sisson, J.C., Sullivan, W., Karess, R. (2004). Reassessing the role and dynamics of nonmuscle myosin II during furrow formation in early Drosophila embryos. Molecular biology of the cell 15, 838–850.

Royou, A., Sullivan, W., Karess, R. (2002). Cortical recruitment of nonmuscle myosin II in early syncytial Drosophila embryos: its role in nuclear axial expansion and its regulation by Cdc2 activity. The Journal of cell biology 158, 127–137.

Rozbicki, E., Chuai, M., Karjalainen, A.I., Song, F., Sang, H.M., Martin, R., Knolker, H.J., MacDonald, M.P., Weijer, C.J. (2015). Myosin-II-mediated cell shape changes and cell intercalation contribute to primitive streak formation. Nature cell biology 17, 397–408.

Saias, L., Swoger, J., D'Angelo, A., Hayes, P., Colombelli, J., Sharpe, J., Salbreux, G., Solon, J. (2015). Decrease in Cell Volume Generates Contractile Forces Driving Dorsal Closure. Developmental cell 33, 611–621.

Samakovlis, C., Hacohen, N., Manning, G., Sutherland, D.C., Guillemin, K., Krasnow, M.A. (1996). Development of the Drosophila tracheal system occurs by a series of morphologically distinct but genetically coupled branching events. Development 122, 1395–1407.

Serra-Picamal, X., Conte, V., Vincent, R., Anon, E., Tambe, D.T., Bazellieres, E., Butler, J.P., Fredberg, J.J., Trepat, X. (2012). Mechanical waves during tissue expansion. Nat Phys 8, 628–U666.

Shiga, Y., Tanaka-Matakatsu, M., Hayashi, S. (1996). A nuclear GFP/β-galactosidase fusion protein as a marker for morphogenesis in living Drosophila. Development, Growth & Differentiation 38, 99–106.

Simoes Sde, M., Blankenship, J.T., Weitz, O., Farrell, D.L., Tamada, M., Fernandez-Gonzalez, R., Zallen, J.A. (2010). Rho-kinase directs Bazooka/Par-3 planar polarity during Drosophila axis elongation. Developmental cell 19, 377–388.

Strutt, D.I., Weber, U., Mlodzik, M. (1997). The role of RhoA in tissue polarity and Frizzled signalling. Nature 387, 292–295.

Tabata, T., Eaton, S., Kornberg, T.B. (1992). The Drosophila hedgehog gene is expressed specifically in posterior compartment cells and is a target of engrailed regulation. Genes & development 6, 2635–2645.

Winter, C.G., Wang, B., Ballew, A., Royou, A., Karess, R., Axelrod, J.D., Luo, L. (2001). Drosophila Rho-associated kinase (Drok) links Frizzled-mediated planar cell polarity signaling to the actin cytoskeleton. Cell 105, 81–91.

Wyatt, T.P.J., Harris, A.R., Lam, M., Cheng, Q., Bellis, J., Dimitracopoulos, A., Kabla, A.J., Charras, G.T., Baum, B. (2015). Emergence of homeostatic epithelial packing and stress dissipation through divisions oriented along the long cell axis. Proceedings of the National Academy of Sciences of the United States of America 112, 5726–5731.

Yamada, S., Nelson, W.J. (2007). Localized zones of Rho and Rac activities drive initiation and expansion of epithelial cell-cell adhesion. The Journal of cell biology 178, 517–527.

Yen, W.W., Williams, M., Periasamy, A., Conaway, M., Burdsal, C., Keller, R., Lu, X., Sutherland, A. (2009). PTK7 is essential for polarized cell motility and convergent extension during mouse gastrulation. Development 136, 2039–2048.

Zallen, J.A., Wieschaus, E. (2004). Patterned gene expression directs bipolar planar polarity in Drosophila. Developmental cell 6, 343–355.

Zecca, M., Struhl, G. (2007). Recruitment of cells into the Drosophila wing primordium by a feed-forward circuit of vestigial autoregulation. Development 134, 3001–3010.

